## Supplementary Figures for "Sex-specific blood-brain barrier alterations and vascular biomarkers underlie chronic stress responses in mice and human depression"

### Extended Figures

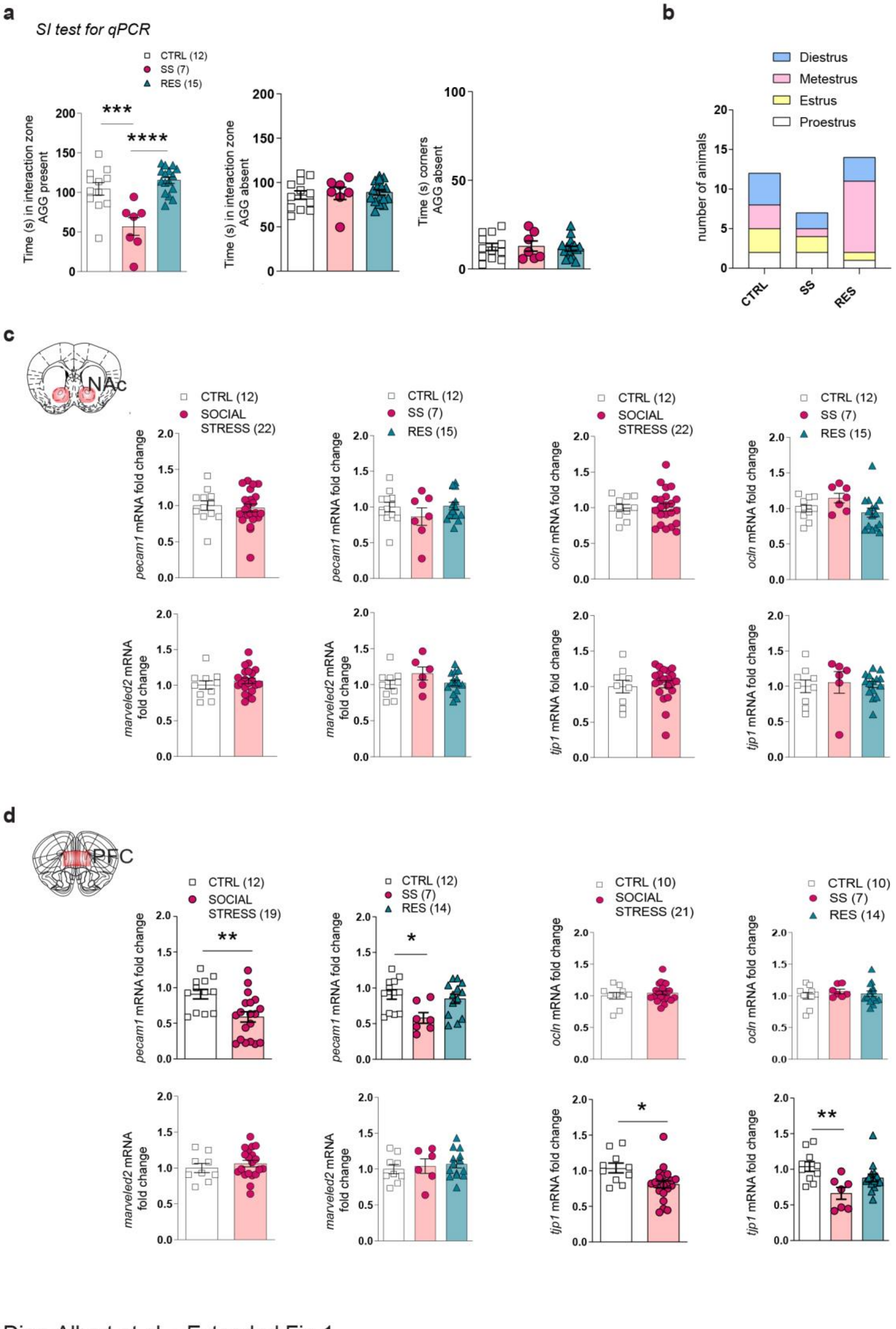


Dion-Albert et al. - Extended Fig.1

**Extended Figure 1. Behavioral phenotype of quantitative PCR (qPCR) experiments and *cldn5* expression in the nucleus accumbens (NAc) and prefrontal cortex (PFC) of female mice after chronic social defeat stress (CSDS). a,** Stress susceptible (SS) mice spent less time in the interaction zone (*****p*<0.0001), when the social target (aggressor, AGG) was present compared to unstressed control (CTRL) and resilient (RES) animals. No significance was observed for time in the interaction zone or in the corners when the social target is absent. **b**, No significant trend was observed between phenotype and phase of the estrus cycle at the time of tissue collection. **c,** Following 10-d CSDS, no difference was observed in *pecam1, ocln, marveled2 or tjp1* mRNA expression in the NAc of female mice, (**d**) while decreased expression of *pecam1* (***p*=0.0048) and *tjp1* (**p*=0.0114) mRNA levels were observed in the PFC of stressed mice. Data represent mean ± s.e.m; number of animals or subjects (*n*) is indicated on graphs. 2-group comparisons were evaluated with unpaired t-tests and one-way ANOVA followed by Bonferroni's multiple comparison test for other graphs. **p*<0.05; ***p*<0.01; ****p*<0.001, *****p*<0.0001.

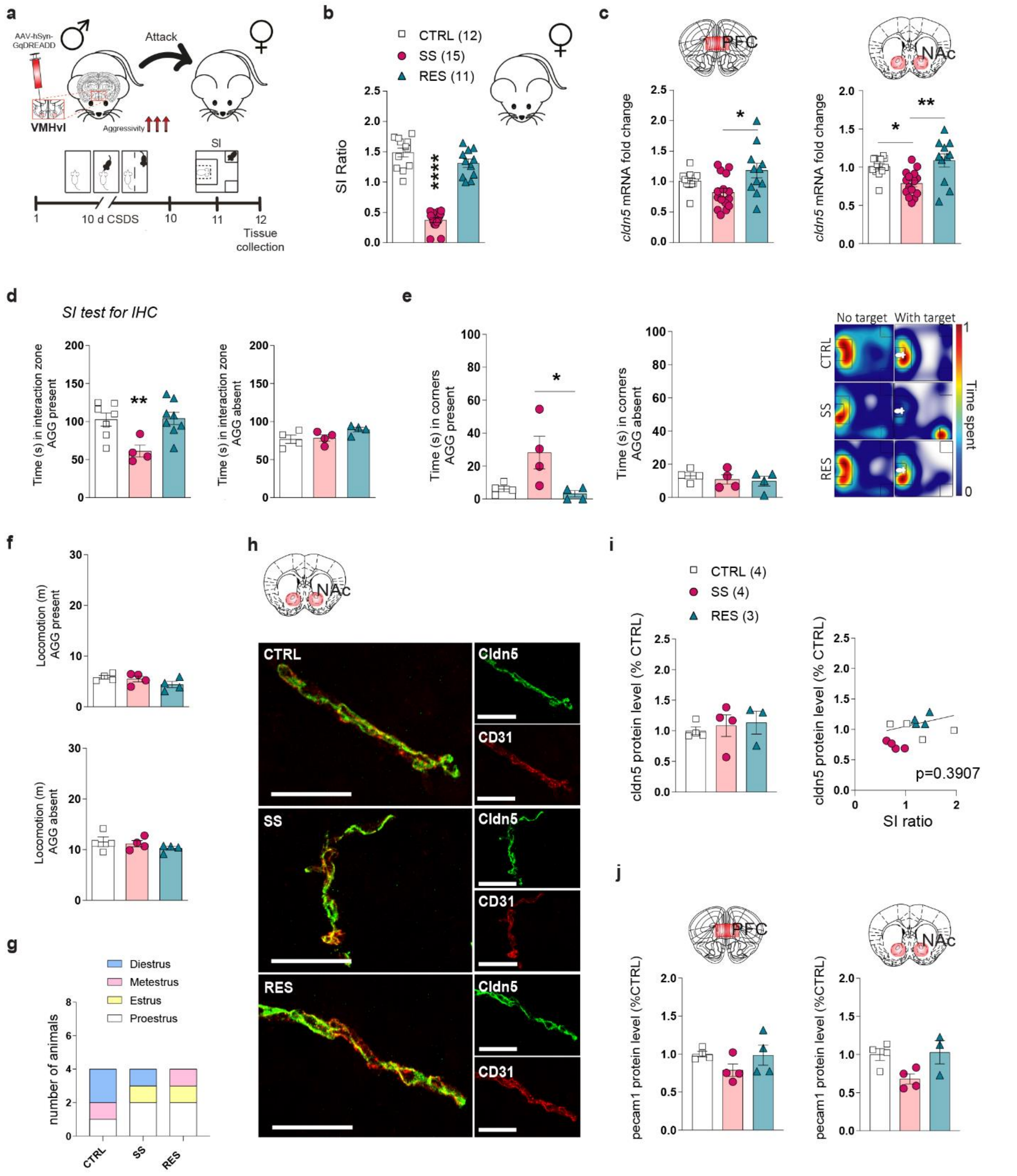


Dion-Albert et al. - Extended Fig.2

**Extended Figure 2. *Cldn5* expression is altered in the nucleus accumbens (NAc) and prefrontal cortex (PFC) of female mice after chronic social defeat stress (CSDS) using**

**hyperaggressive males however no significant change was observed at protein level in the NAc. a**, Schematic representation of the CSDS female paradigm developed by Takahashi et al. (2017). **b,** Like in the conventional male paradigm, 10-d physical exposure to hyperaggressive CD1 males (aggressor, AGG) leads to two female subpopulations – stress-susceptible (SS) and resilient (RES) - as defined by the social interaction (SI) test (****$p<0.0001$). **c,** Stress susceptibility was associated with a loss of *cldn5* expression in the PFC (*$p=0.0122$) and NAc (**$p=0.0013$) of female mice (normalized on *pecam1)* in this CSDS model. **d,** Behavioral phenotyping of the female cohort used for immunohistochemistry with SS mice spending less time in the interaction zone (**$p=0.0013$) and (**e**) increased time in the corners when the social target was present compared to unstressed control (CTRL) and RES animals. No significant difference was observed when the social target was absent. **f,** No difference was observed in total travelled distance (m) either when the social target was present or absent. **g,** No significant trend was observed between phenotype and phase of the estrus cycle at the time of tissue collection. **h,i,** cldn5 protein levels in the NAc were not significantly different between CTRL, SS and RES animals, and do not correlate with social avoidance ($p=0.3907$). Scale bars, 20μm. **j,** No difference was observed in protein levels of endothelial marker Pecam1 in the PFC or NAc of animals. Data represent mean ± s.e.m; number of animals or subjects ($n$) is indicated on graphs. 2-group comparisons were evaluated with unpaired t-tests and one-way ANOVA followed by Bonferroni's multiple comparison test for other graphs. *$p<0.05$; **$p<0.01$.

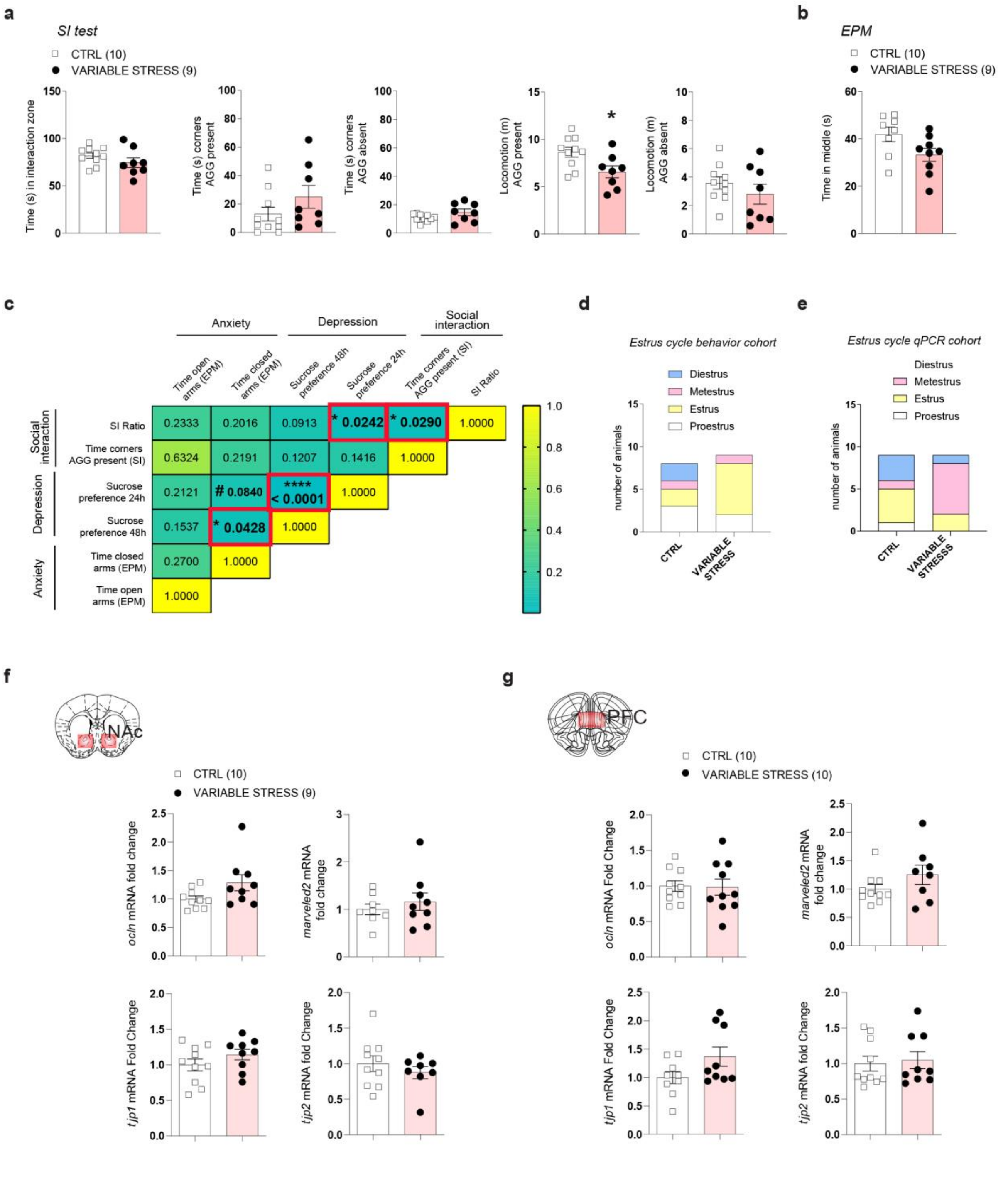


Dion-Albert et al. - Extended Fig.3

**Extended Figure 3. Behavioral phenotypes and quantitative PCR (qPCR) experiments in the nucleus accumbens (NAc) and prefrontal cortex (PFC) of female mice after 6d subchronic variable stress (SCVS). a,** following 6-d SCVS, variable stress and control (CTRL) females spent similar times in the interaction zone and corners when the social target (aggressor, AGG) was present or absent. SCVS females travelled less distance when the aggressor was present (**p*=0.0202) vs CTRL. **b,** No significant difference was observed for time in the middle of the EPM between stressed and CTRL animals. **c,** Intraindividual correlations between various behavioral data points shows association between social interaction, anxiety- and depressive-like behaviors. P values in the boxes refer to the strength of the correlation between behaviors. **d,e,** No significant trend was observed between phenotype and phase of the estrus cycle between CTRL and variable stress animals in the behavior (**d**) and qPCR (**e**) cohorts. **f,g,** Following 6-d SCVS, no difference was observed in *ocln, marveled2, tjp1, or tjp2* mRNA levels in the NAc (**f**) or PFC (**g**) of stressed female mice. Data represent mean ± s.e.m; number of animals or subjects (*n*) is indicated on graphs. 2-group comparisons were evaluated with unpaired t-tests and one-way ANOVA followed by Bonferroni's multiple comparison test for other graphs. #*p*<0.1; **p*<0.05; *****p*<0.0001.

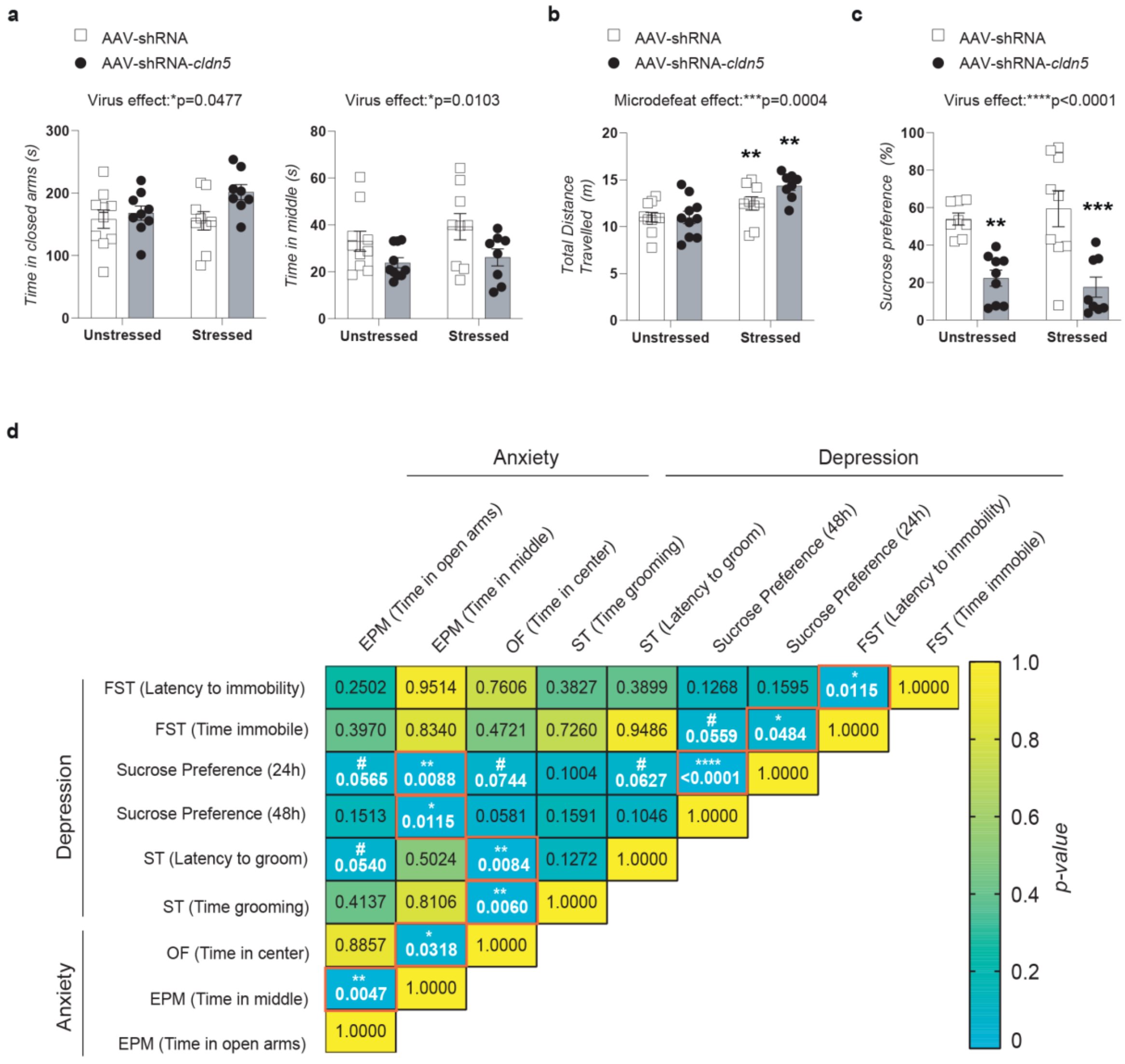


Dion-Albert et al. - Extended Fig.4

**Extended Figure 4. Supplementary behavior results for AAV-shRNA-*cldn5* injected mice. a,** A main virus effect was revealed in AAV-shRNA vs AAV-shRNA-*cldn5* injected mice for time in closed arms (**p*=0.0477) and middle area (**p* =0.0103) in the elevated plus maze (EPM) test. **b,** Animals who underwent subthreshold microdefeat (stressed mice) travelled significantly higher distance than their unstressed counterparts (****p*=0.0004). **c**, a highly significant main virus effect was found at 48h in the sucrose preference test (*****p*<0.0001). **d,** Intraindividual correlations between various behavioral data points shows strong association between social interaction, anxiety- and depressive-like behavior between AAV-shRNA and AAV-shRNA-*cldn5* animals. P values in the boxes refer to the strength of the correlation between behaviors. Data represent mean ± s.e.m; number of animals or subjects (*n*). Correlations were evaluated with Pearson's correlation coefficient and two-way ANOVA followed by Bonferroni's multiple comparison test for other graphs. #$p<0.1$; *$p<0.05$; **$p<0.01$; ***$p<0.001$.

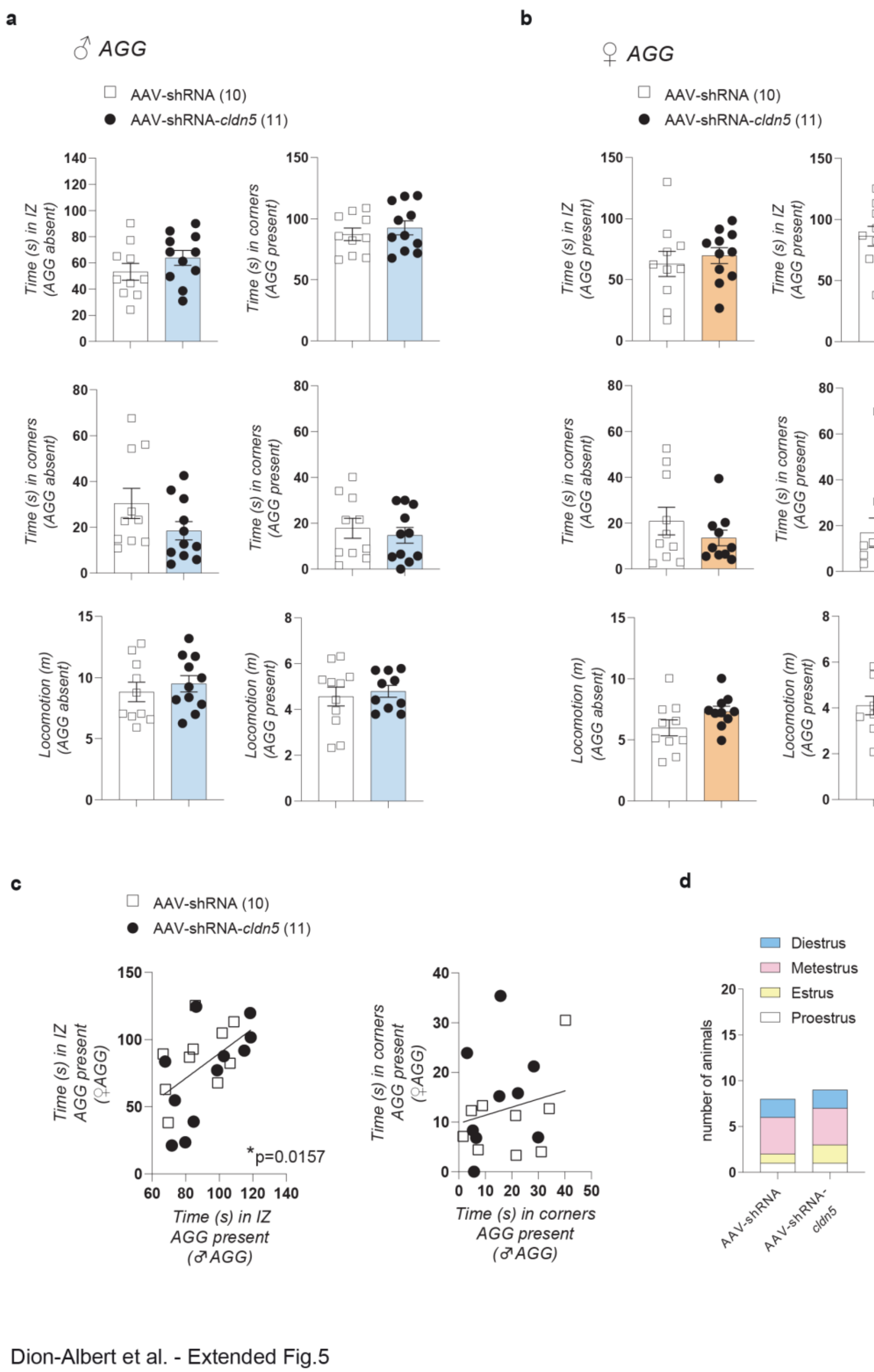


**Extended Figure 5. Supplementary SI behavior results for AAV-shRNA and AAV-shRNA-*cldn5* injected mice. a,b,** Time in the interaction zone (IZ), corners and total distance travelled was not significantly different between AAV-shRNA and AAV-shRNA-*cldn5*-injected mice, whether the male (**a**) or female (**b**) social target (aggressor, AGG) was present or absent. **c**, Time in the IZ when the female AGG was present significantly correlated to the time in the IZ when the male AGG was present (**p*=0.0157), but no correlation was found for time in corners. **d,** No significant trend was observed between phenotype and phase of the estrus cycle between AAV-shRNA and AAV-shRNA-*cldn5* injected animal. Data represent mean ± s.e.m; number of animals or subjects (*n*) is indicated on graphs. Correlations were evaluated with Pearson's correlation coefficient and 2-group comparisons were evaluated with unpaired t-tests and one-way ANOVA followed by Bonferroni's multiple comparison test for other graphs. *$p$<0.05

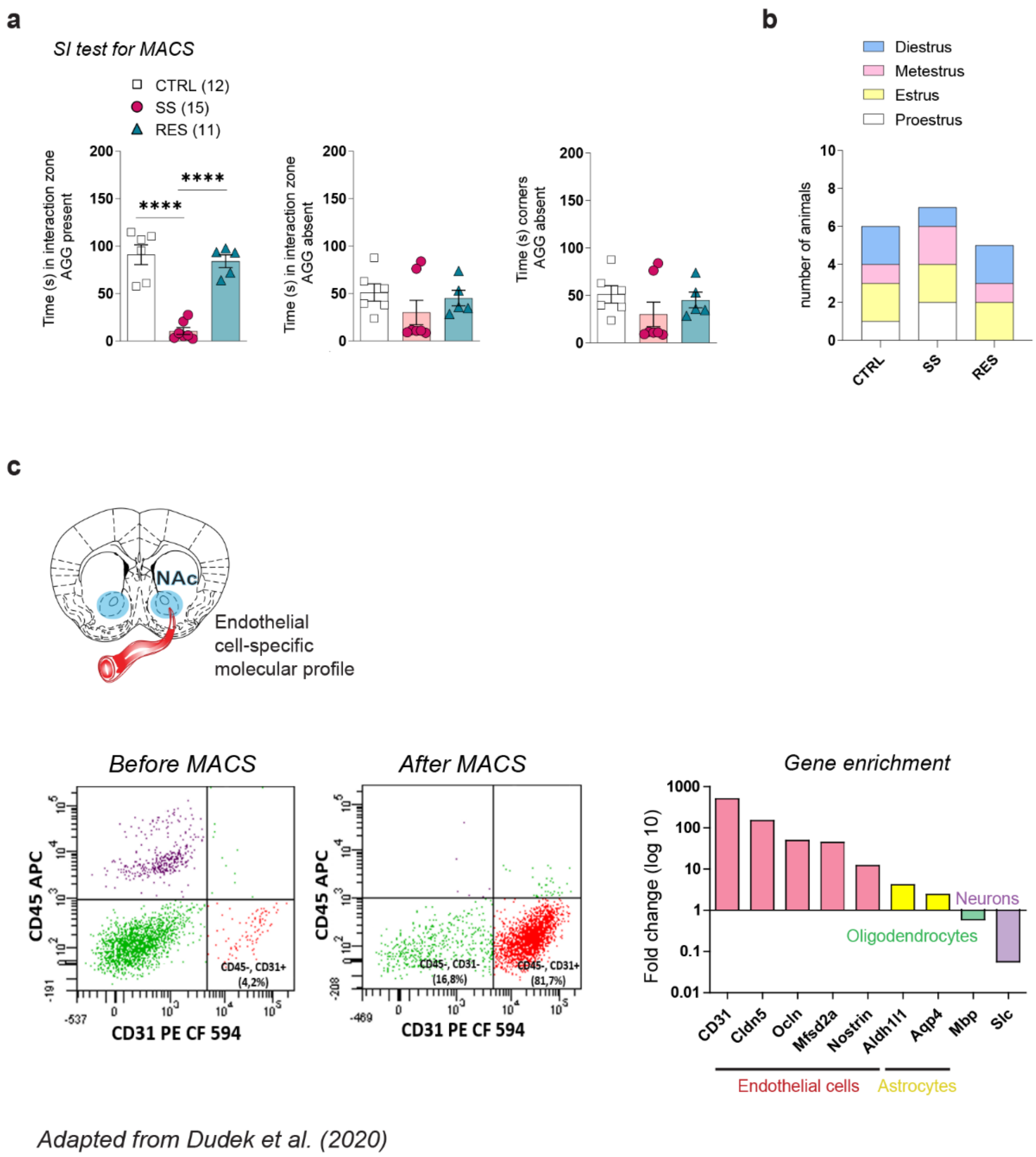


Dion-Albert et al. - Extended Fig.6

**Extended Figure 6. Behavioral phenotype of CSDS for transcriptomic experiment of PFC endothelial cells and validation of magnetic-activated cell sorting (MACS). a,** Stress susceptible (SS) mice spent less time in the interaction zone, when the social target (aggressor, AGG) was present compared to unstressed control (CTRL) and resilient (RES) animals (*****p*<0.0001). No significance was observed for time in the interaction zone or in the corners when the social target is absent. **b**, No significant trend was observed between phenotype and phase of the estrus cycle at the time of tissue collection. **c**, Enrichment of endothelial cells following MACS was confirmed using flow cytometry and qPCRs (adapted from Dudek et al.[8]). As shown on the left, CD31+ cell population increases from ~4% in the heterogenous preparation to >80% after MACS. Endothelial cell-specific gene expression (CD31*;* claudin-5, *cldn5;* occludin, *ocln;* major facilitator superfamily domain-containing protein 2, *mfsd2a; nostrin*) is highly increased as well when compared to genes related to astrocytes (aldehyde dehydrogenase 1 family member l1, *aldh1l1;* aquaporin 4, *aqp4*), oligodendrocytes (myelin basic protein, *mbp*) or neurons (solute carrier, *slc*). Data represent mean ± s.e.m; number of animals or subjects (*n*) is indicated on graphs. 3-group comparisons were evaluated with one-way ANOVA followed by Bonferroni's multiple comparison tests. ****$p<0.0001$.

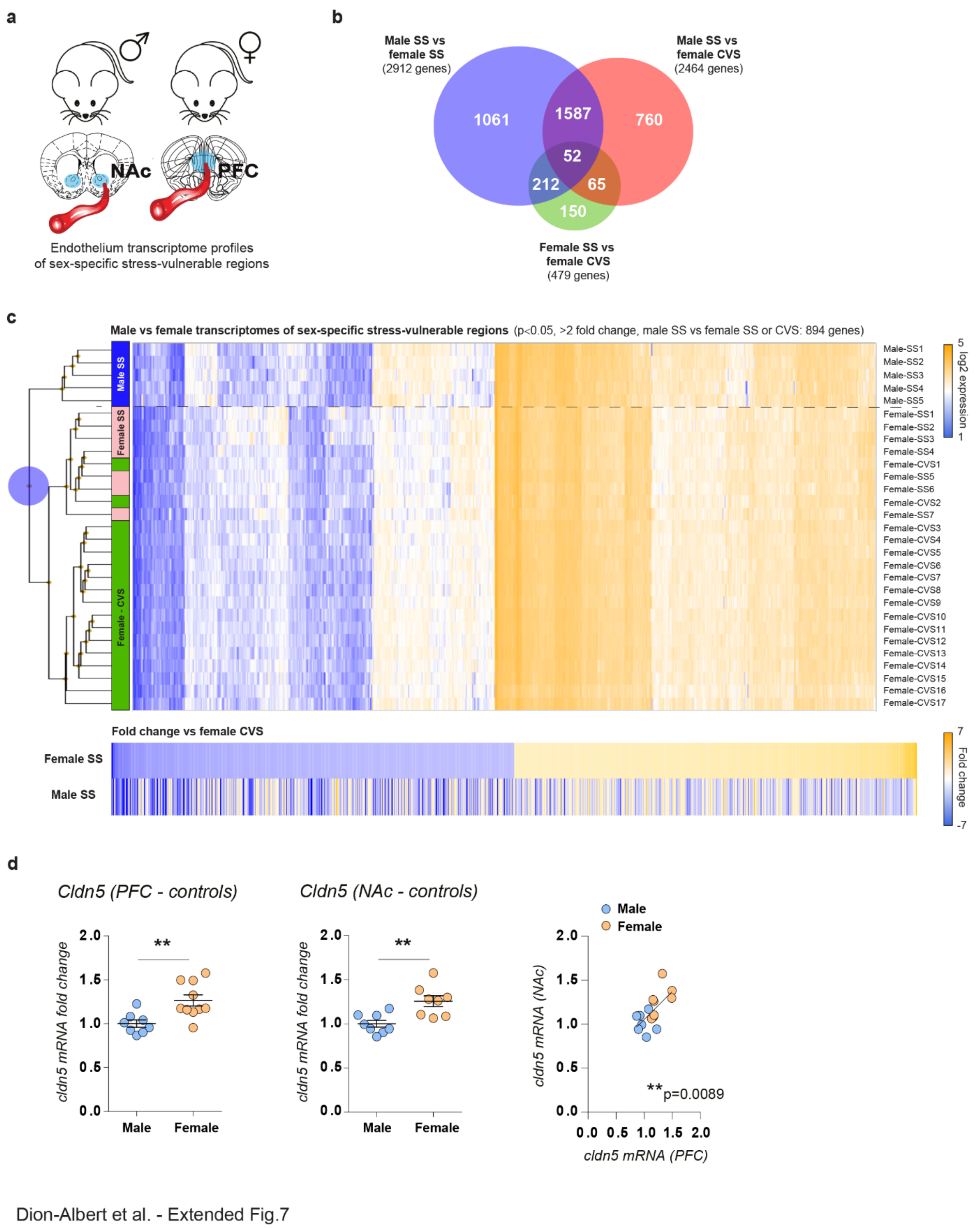


**Extended Figure 7. Comparison of stress-induced transcriptomic changes in endothelial cells in male vs female's most vulnerable brain region. a,** Schematic outline of sex- and regionspecific comparison. **b,** Venn diagrams reveals poor overlap of gene expression changes when group comparisons were performed, with stress-susceptible (SS) male vs SS female showing the highest number of differentially regulated genes. **c,** Hierarchical clustering reveals more genes regulated in the same direction between females exposed to different stress paradigms then male vs females exposed to CSDS (SS only). Significance was set at p<0,05 and ±2-fold change from the mean. **d,** Stress-naïve female mice exhibit higher *cldn5* mRNA levels in the prefrontal cortex (PFC) (***p*=0.0044) and nucleus accumbens (NAc) (***p*=0.0034) than males, which are positively correlated with each other (***p*=0.0089). Data represent mean ± s.e.m; number of animals or subjects (*n*) is indicated on graphs. Correlations were evaluated with Pearson's correlation coefficient and 2-group comparisons were evaluated with unpaired t-tests. ** *p*<0.01

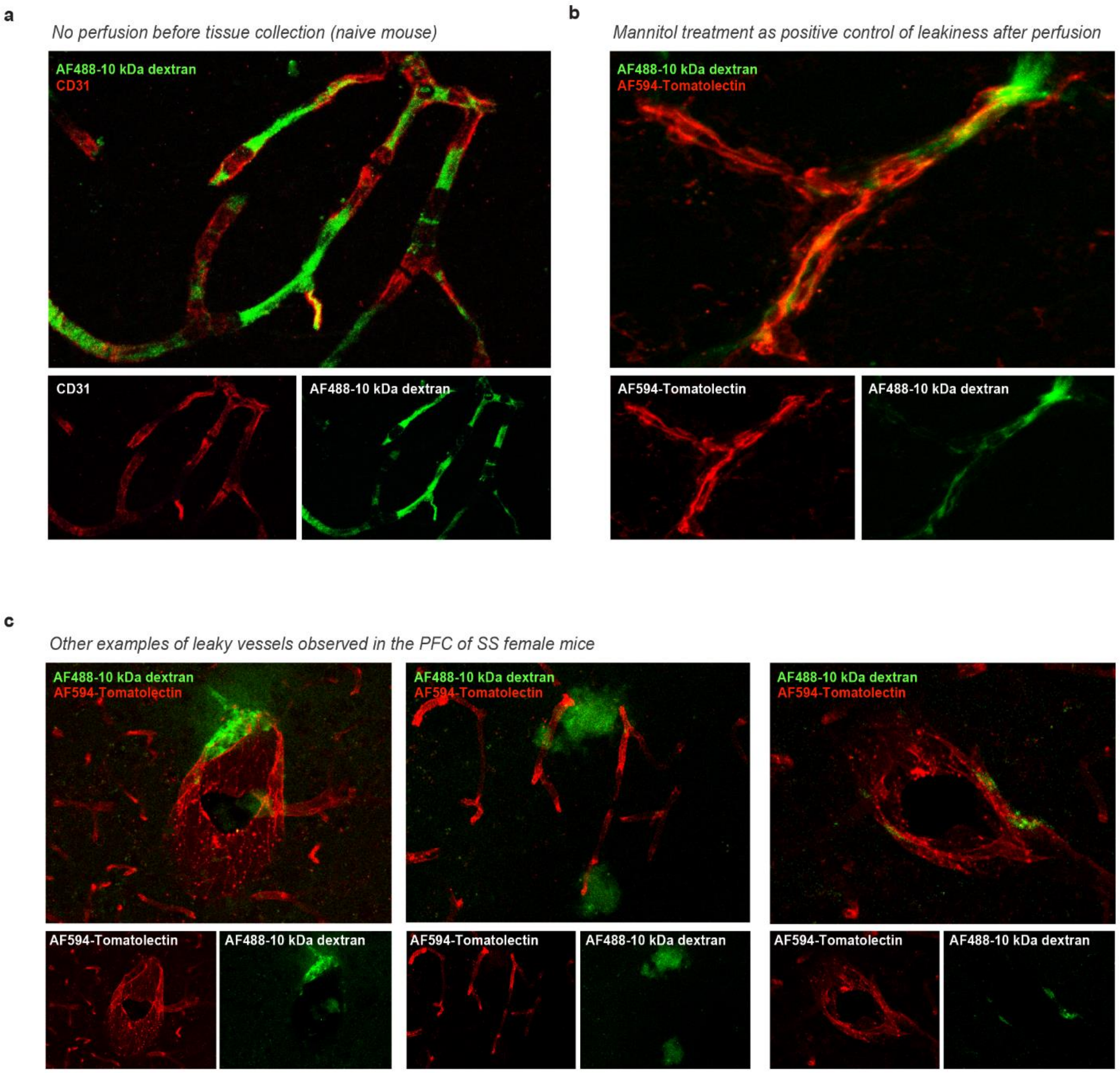


Dion-Albert et al. - Extended Fig.8

**Extended Figure 8. Supplementary images for blood vessel leakiness in the prefrontal (PFC) of female mice. a,** Dextran AlexaFluor488 (AF488-dextran) could be detected in only PFC blood vessels 30 minutes after retro-orbital injection. **b,** AF488-dextran extravasation was observed after 30-min circulation and intravenous administration of mannitol for 5 min prior perfusion, as a positive control. **c,** Additional images of PFC BBB leakiness in SS mice following CSDS, showing dye accumulation in the parenchyma and perivascular space of blood vessels.

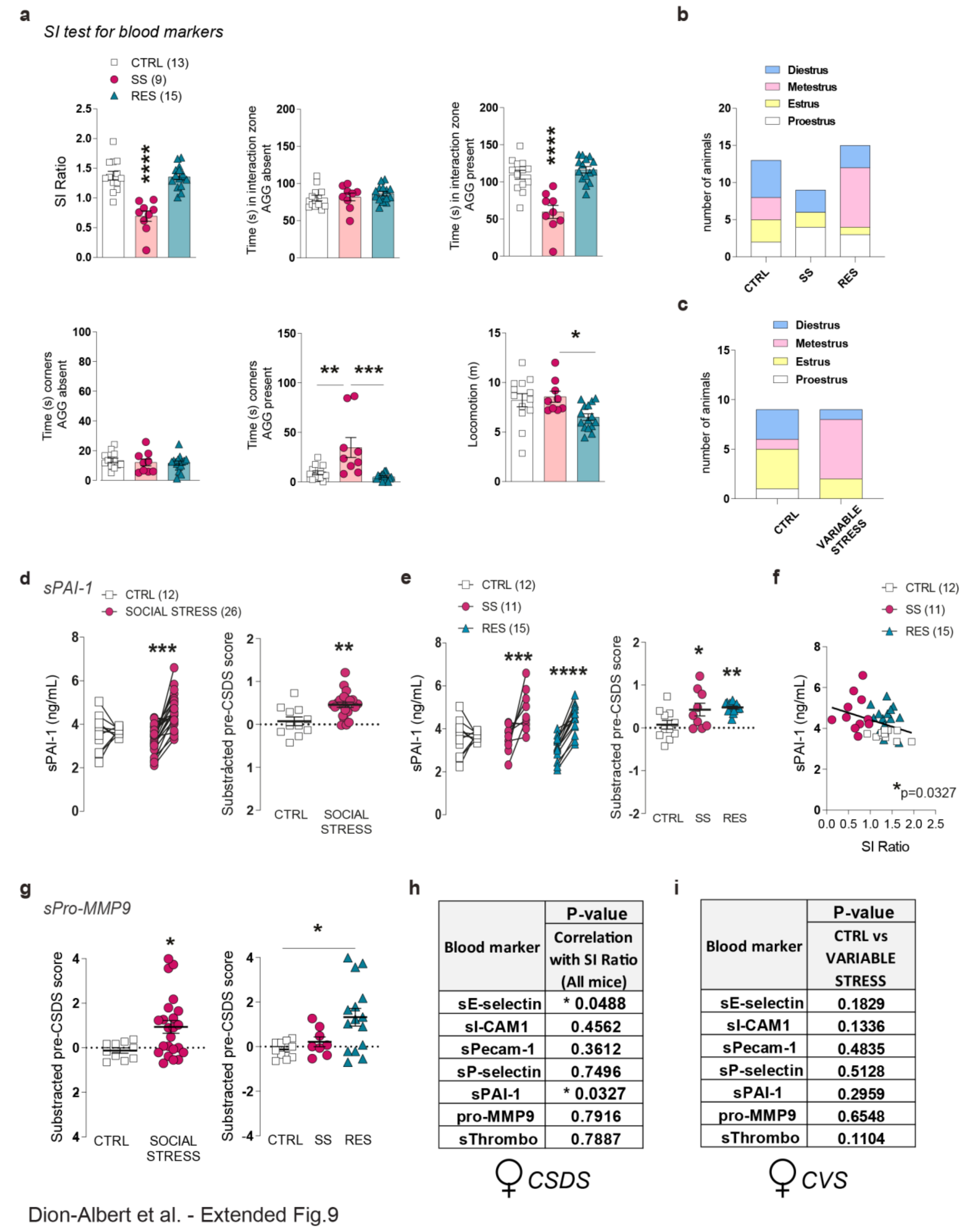


**h**

| Blood marker | P-value Correlation with SI Ratio (All mice) |
|---|---|
| sE-selectin | * 0.0488 |
| sI-CAM1 | 0.4562 |
| sPecam-1 | 0.3612 |
| sP-selectin | 0.7496 |
| sPAI-1 | * 0.0327 |
| pro-MMP9 | 0.7916 |
| sThrombo | 0.7887 |

**i**

| Blood marker | P-value CTRL vs VARIABLE STRESS |
|---|---|
| sE-selectin | 0.1829 |
| sI-CAM1 | 0.1336 |
| sPecam-1 | 0.4835 |
| sP-selectin | 0.5128 |
| sPAI-1 | 0.2959 |
| pro-MMP9 | 0.6548 |
| sThrombo | 0.1104 |

**Extended Figure 9. Behavioral phenotype and supplemental data for Milliplex experiments in female mice. a,** Stress-susceptible (SS) mice spent less time in the interaction zone (*****p*<0.0001) and increased time in the corners (****p*=0.0002) when the social target (aggressor, AGG) was present compared to unstressed control (CTRL) and resilient (RES) animals. No significant difference was observed when the social target was absent. RES animal travelled less distance (m) than SS mice in total. **b,** No significant trend was observed between phenotype and phase of the estrus cycle at the time of tissue collection for the chronic social defeat stress (CSDS) cohort (**b**) or the subchronic variable stress (SCVS) cohort (**c**). **d-f,** soluble plasminogen activator inhibitor 1 (sPAI-1) levels (ng/mL) were significantly increased in social stress (****p*=0.0004, pre-CSDS score ***p*=0.0016), SS (***p*=0.003, pre-CSDS score **p*=0.0478) and RES (*****p*<0.0001, pre-CSDS score ***p*=0.0081) mice when compared to their baseline pre-CSDS levels and correlated with social avoidance (**p*=0.0327). **g, s**Pro-MMP9 levels were significantly increased in social stress (**p*=0.0245) and RES (*******p*=0.0077) animals following CSDS, when compared to their individual baseline levels. **h**, Individual correlation between serum levels of circulating vascular markers and SI Ratio was significant only for sE-selectin (**p*=0.0488) and sPAI-1 (**p*=0.0327) following CSDS. **i**, Circulating levels of vascular biomarkers following 6-d SCVS were not significantly different when compared to CTRL. Data represent mean ± s.e.m; number of animals or subjects (*n*) is indicated on graphs. 2-group comparisons were evaluated with unpaired t-tests and one-way ANOVA followed by Bonferroni's multiple comparison test for other graphs. Correlations were evaluated with Pearson's correlation coefficient and 2-group comparisons were evaluated with unpaired t-tests. **p*<0.05; ***p*<0.01; ****p*<0.001; *****p*<0.0001

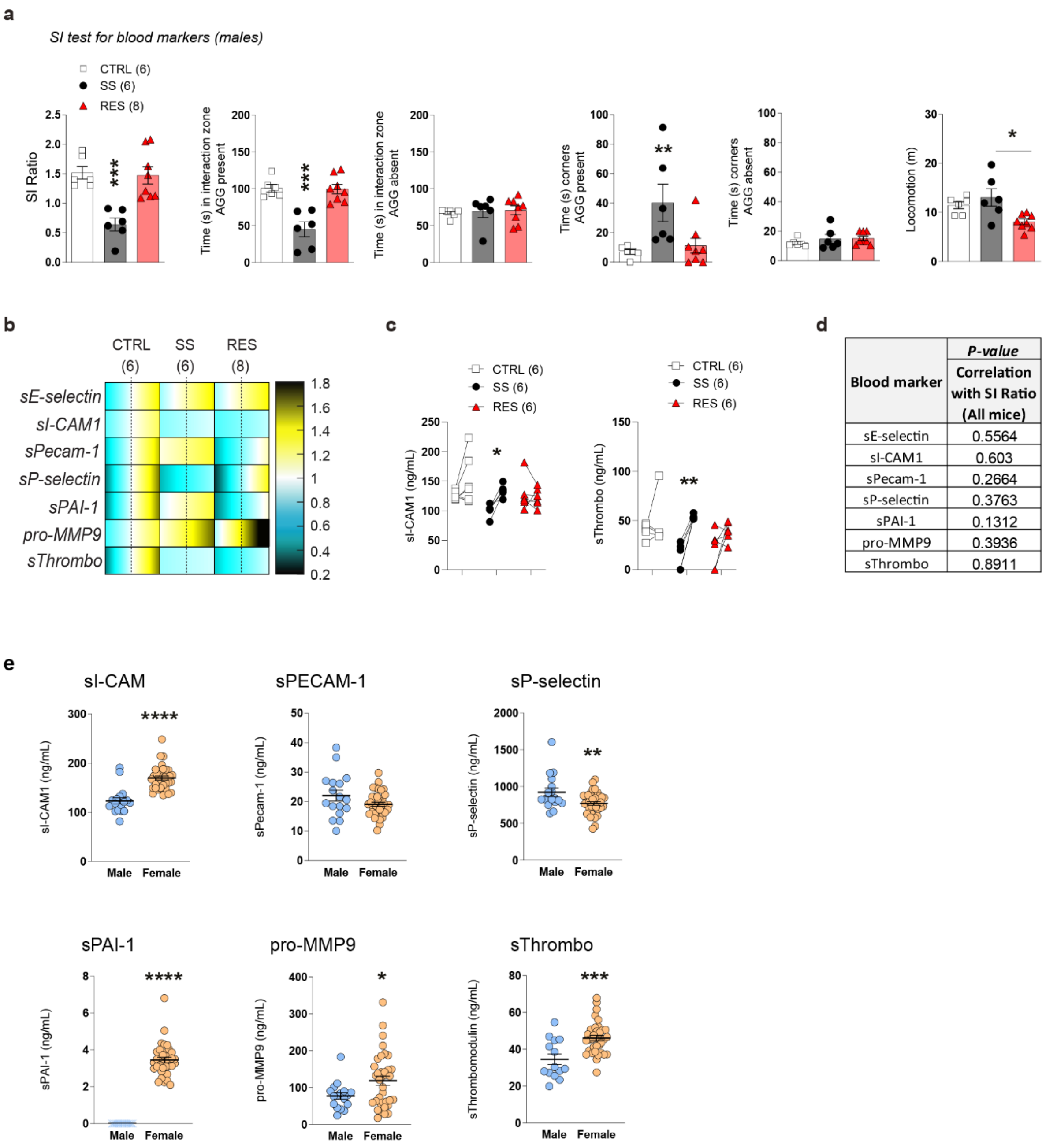


Dion-Albert et al. - Extended Fig.10

**Extended Figure 10. Behavioral phenotype and supplemental data for Milliplex experiments in male mice and baseline sex-differences in circulating vascular markers. a,** Stress-susceptible (SS) mice spent less time in the interaction zone (****p*=0.0003) and increased time in the corners (*****p*<0.0001) when the social target (aggressor, AGG) was present compared to unstressed control (CTRL) and resilient (RES) animals. No significant difference was observed when the social target was absent. RES animal travelled less distance (m) than SS mice in total. **b,** Heatmap of circulating blood markers normalized to CTRL after 10-d chronic social defeat stress (CSDS). **c,** sI-CAM1 and sThrombomodulin levels (ng/mL) significantly increased only in SS mice, when compared to their individual baseline levels. **d,** None of the investigated markers correlated to social avoidance in male mice. **e,** Baseline sex-differences in circulating sI-CAM1 (****$p$<0.0001), sPECAM-1 ($p$=0.0802), sP-selectin (**$p$=0.0066), sPAI-1 (****$p$<0.0001), sPro-MMP9 (*$p$=0.0346) and sThrombomodulin (***$p$=0.0003) levels. Data represent mean ± s.e.m; number of animals or subjects (*n*) is indicated on graphs. 2-group comparisons were evaluated with unpaired t-tests and one-way ANOVA followed by Bonferroni's multiple comparison test for other graph. *$p$<0.05; **$p$<0.01; ***$p$<0.001; ****$p$<0.0001.
