## Supplementary tables for "Sex-specific blood-brain barrier alterations and vascular biomarkers underlie chronic stress responses in mice and human depression"

**Table 1. Mouse Primers qPCR**

| Gene | Ref Seq # | Assay ID | Forward primer | Reverse primer |
|---|---|---|---|---|
| *pecam1* | NM_008816(2) | Mm.PT.58.43167370 | PrimeTime®qPCR primers Exon Location 7-8 | |
| *cdh5* | NM_009868(1) | Mm.PT.58.8747496 | PrimeTime®qPCR primers Exon Location 7-8 | |
| *fgf2* | NM_008006(1) | Mm.PT.56a.5129235 | PrimeTime®qPCR primers Exon Location 1-3 | |
| *angpt2* | NM_007426(1) | Mm.PT.58.29139310 | PrimeTime®qPCR primers Exon Location 1a-2 | |
| *vegfa* | NM_009505(3) | Mm.PT.58.14200306 | PrimeTime®qPCR primers Exon Location 1-2 | |
| *cldn1* | NM_016674(1) | Mm.PT.58.41579102 | PrimeTime®qPCR primers Exon Location 2-3 | |
| *cldn3* | NM_009902(1) | Mm.PT.58.43310459.g | PrimeTime®qPCR primers Exon Location 1-1 | |
| *cldn5* | NM_013805(1) | 5'-TTTCTTCTATGCGCAGTTGG-3' | 5'-GCAGTTTGGTGCCTACTTCA-3' | |
| *cldn12* | NM_001193659(3) | Mm.PT.58.41535303 | PrimeTime®qPCR primers Exon Location 3a-5 | |
| *tjp1* | NM_001163574(2) | Mm.PT.58.29459730 | PrimeTime®qPCR primers Exon Location 18-19 | |
| *tjp2* | NM_001198985(2) | Mm.PT.58.16834535 | PrimeTime®qPCR primers Exon Location 22-23 | |
| *tjp3* | NM_013769(1) | Mm.PT.58.43961106 | PrimeTime®qPCR primers Exon Location 15-17 | |
| *ocln* | NM_008756(1) | Mm.PT.58.42749240 | PrimeTime®qPCR primers Exon Location 7-9 | |
| *marveld2* | NM_001038602(2) | Mm.PT.58.7719303 | PrimeTime®qPCR primers Exon Location 5-7 | |
| *mfsd2a* | NM_029662(1) | Mm.PT.58.32675283 | PrimeTime®qPCR primers Exon Location 13-14 | |
| *ntn1* | NM_008744(1) | Mm.PT.58.41162316 | PrimeTime®qPCR primers Exon Location 3-4 | |
| *gapdh* | NM_008084(1) | Mm.PT.39a.1 | PrimeTime®qPCR primers Exon Location 2-3 | |
| *nostrin* | NM_181547(1) | Mm.PT.58.12259437 | PrimeTime®qPCR primers Exon Location 14-16 | |
| *aldh1l1* | NM_027406(1) | Mm.PT.58.7775479 | PrimeTime®qPCR primers Exon Location 13-15 | |
| *aqp4* | NM_009700(1) | Mm.PT.58.9080805 | PrimeTime®qPCR primers Exon Location 1-2 | |
| *mbp* | NM_001025254 (4) | Mm.PT.58.28532164 | PrimeTime®qPCR primers Exon Location 6-9 | |
| *slc17a6* | Mbp NM_080853(1) | Mm.PT.58.10363705 | PrimeTime®qPCR primers Exon Location 9-10 | |

**Table 2. Human Primers qPCR**

| Gene | Ref Seq # | Assay ID | Primers |
|---|---|---|---|
| *CLDN5* | NM_003277(2) | Hs.PT.58.1483777.g | PrimeTime®qPCR primers Exon Location 1-1 |
| *GAPDH* | NM_002046(1) | Hs.PT.39a.22214836 | PrimeTime®qPCR primers Exon Location 2-3 |

**Table 3. Complete demographic for human brain cohort**

| Gender | Age | Post mortem interval | Cause of death | Depressive symptoms | History of abuse | Antidepressant treatment |
|---|---|---|---|---|---|---|
| F | 66 | 61 | Accident | No | N/A | No |
| F | 72 | 17 | Natural | No | No | No |
| F | 45 | 44 | Suicide | No | Yes | Yes |
| F | 81 | 13,5 | Natural | No | N/A | No |
| F | 78 | 7,5 | Accident | No | Yes | No |
| F | 76 | 7 | Accident | No | No | Yes |
| F | 68 | 8,5 | Natural | No | N/A | No |
| F | 85 | 12,0 | Accident | No | N/A | No |
| F | 67 | 16,6 | Accident | No | N/A | No |
| F | 57 | 39 | Accident | No | No | Yes |
| F | 51 | 111 | Natural | No | No | Yes |
| F | 28 | 80 | Undetermined | No | No | No |
| M | 47 | 12 | Natural | No | No | No |
| M | 41 | 24 | Natural | No | No | No |
| M | 30 | 30 | Accident | No | No | No |
| M | 19 | 27.75 | Suicide | No | N/A | No |
| M | 46 | 19.5 | Natural | No | No | No |
| M | 32 | 29.5 | Accident | No | N/A | No |
| M | 33 | 18 | Suicide | No | Yes | No |
| M | 42 | 63 | Accident | No | N/A | No |
| M | 55 | 24 | Accident | No | Yes | No |
| M | 46 | 59 | Suicide | No | N/A | No |
| M | 52 | 10 | Natural | No | N/A | No |
| M | 29 | 4 | Suicide | No | N/A | No |
| M | 59 | 23.5 | Accident | No | N/A | No |
| F | 25 | 20 | Suicide | Yes | N/A | No |
| F | 55 | 36 | Suicide | Yes | No | No |
| F | 25 | 56 | Suicide | Yes | N/A | No |
| F | 32 | 41 | Suicide | Yes | N/A | No |
| F | 55 | 2,5 | Suicide | Yes | Yes | No |
| F | 44 | 60 | Suicide | Yes | N/A | No |
| F | 40 | 106,5 | Natural | Yes | N/A | No |
| F | 52 | 71,42 | Suicide | Yes | N/A | No |
| F | 46 | 58,43 | Suicide | Yes | N/A | No |
| F | 52 | 70,00 | Suicide | Yes | N/A | No |
| F | 46 | 15 | Suicide | Yes | N/A | Yes |
| F | 40 | 49,5 | Suicide | Yes | N/A | Yes |
| F | 54 | 28,5 | Suicide | Yes | Yes | Yes |

| | | | | | | |
|---|---|---|---|---|---|---|
| F | 49 | 14,5 | Suicide | Yes | N/A | Yes |
| F | 36 | 7,5 | Suicide | Yes | No | Yes |
| F | 55 | 36 | Suicide | Yes | N/A | Yes |
| F | 59 | 3 | Suicide | Yes | No | Yes |
| F | 55 | 70,72 | Suicide | Yes | N/A | Yes |
| F | 56 | 74,78 | Accidental | Yes | N/A | Yes |
| F | 42 | 59,20 | Suicide | Yes | N/A | Yes |
| F | 56 | 78,00 | Natural | Yes | N/A | Yes |
| F | 60 | 74,00 | Suicide | Yes | N/A | Yes |
| F | 72 | 44,00 | Suicide | Yes | N/A | Yes |
| M | 49 | 2.5 | Suicide | Yes | No | No |
| M | 53 | 33.5 | Suicide | Yes | Yes | No |
| M | 38 | 30 | Suicide | Yes | N/A | No |
| M | 28 | 36 | Suicide | Yes | N/A | No |
| M | 29 | 27 | Suicide | Yes | No | No |
| M | 68 | 43 | Suicide | Yes | N/A | No |
| M | 63 | 50 | Suicide | Yes | N/A | No |
| M | 48 | 49 | Suicide | Yes | No | No |
| M | 67 | 56 | Suicide | Yes | N/A | No |
| M | 52 | 29 | Suicide | Yes | N/A | No |
| M | 53 | 14 | Suicide | Yes | N/A | No |
| M | 51 | 54 | Suicide | Yes | N/A | No |
| M | 39 | 25.5 | Suicide | Yes | N/A | No |
| M | 49 | 32 | Suicide | Yes | N/A | No |
| M | 40 | 22 | Suicide | Yes | N/A | No |

**Table 4. Complete demographic data for human serum cohort**

| Gender | Age | Depressive symptoms | Current suicidal thoughts | History of abuse |
|---|---|---|---|---|
| F | 28 | No | No | No |
| F | 29 | No | No | No |
| F | 22 | No | No | No |
| F | 20 | No | No | No |
| F | 29 | No | No | No |
| F | 21 | No | No | No |
| F | 25 | No | No | No |
| F | 29 | No | No | No |
| F | 22 | No | No | No |
| F | 26 | No | No | Yes |
| F | 20 | No | No | No |
| F | 22 | No | No | No |
| F | 30 | No | No | Yes |
| F | 29 | No | No | Yes |
| F | 22 | No | No | No |
| F | 18 | No | Yes | Yes |
| F | 18 | No | No | Yes |
| F | 20 | No | No | No |
| F | 20 | No | No | No |
| F | 19 | No | No | No |
| F | 18 | No | No | No |
| F | 21 | No | No | No |
| F | 29 | Yes | No | No |
| F | 26 | Yes | No | No |
| F | 24 | Yes | No | No |
| F | 21 | Yes | Yes | Yes |
| F | 25 | Yes | Yes | Yes |
| F | 21 | Yes | Yes | No |
| F | 27 | Yes | Yes | Yes |
| F | 23 | Yes | Yes | No |
| F | 24 | Yes | No | No |
| F | 18 | Yes | N/A | No |
| F | 20 | Yes | N/A | Yes |
| F | 29 | Yes | N/A | Yes |
| F | 29 | Yes | N/A | Yes |
| F | 21 | Yes | N/A | Yes |
| F | 19 | Yes | N/A | Yes |
| F | 27 | Yes | N/A | Yes |

| | | | | |
|---|---|---|---|---|
| F | 26 | Yes | N/A | Yes |
| F | 23 | Yes | N/A | Yes |
| F | 21 | Yes | No | No |
| F | 23 | Yes | Yes | Yes |
| F | 25 | Yes | Yes | Yes |
| F | 24 | Yes | Yes | No |
| F | 25 | Yes | No | No |
| F | 22 | Yes | No | Yes |
| F | 24 | Yes | No | Yes |
| F | 28 | Yes | No | Yes |
| F | 30 | Yes | No | Yes |
| F | 30 | Yes | Yes | Yes |
| M | 27 | No | No | Y |
| M | 26 | No | No | N |
| M | 25 | No | No | N |
| M | 28 | No | No | Y |
| M | 25 | No | No | N |
| M | 27 | No | No | N |
| M | 28 | No | No | N |
| M | 24 | No | No | Y |
| M | 27 | No | No | N |
| M | 27 | No | No | N |
| M | 30 | No | No | N |
| M | 24 | No | No | N |
| M | 19 | Yes | No | N |
| M | 23 | Yes | Yes | Y |
| M | 30 | Yes | Yes | Y |
| M | 26 | Yes | Yes | Y |
| M | 18 | Yes | No | N |
| M | 23 | Yes | Yes | Y |
| M | 25 | Yes | No | Y |
| M | 30 | Yes | Yes | Y |
| M | 24 | Yes | No | Y |
| M | 24 | Yes | Yes | N |
| M | 27 | Yes | Yes | N |
| M | 26 | Yes | No | Y |
